# Interrupting T cell memory ameliorates exaggerated metabolic response to weight cycling

**DOI:** 10.1101/2025.01.17.633599

**Authors:** Jamie N. Garcia, Matthew A. Cottam, Alec S. Rodriguez, Anwar F. Hussein Agha, Nathan C. Winn, Alyssa H. Hasty

## Abstract

People frequently experience cycles of weight gain and loss. This weight cycling has been demonstrated, in humans and animal models, to increase cardiometabolic disease and disrupt glucose homeostasis. Obesity itself — and to an even greater extent weight regain — causes adipose tissue inflammation, resulting in metabolic dysfunction. Studies show that even after weight loss, increased numbers of lipid associated macrophages and memory T cells persist in adipose tissue and become more inflammatory upon weight regain. These findings suggest that the immune system retains a "memory" of obesity, which may contribute to the elevated inflammation and metabolic dysfunction associated with weight cycling. Here, we show that blocking the CD70-CD27 axis, critical for formation of immunological memory, decreases the number of memory T cells and reduces T cell clonality within adipose tissue after weight loss and weight cycling. Furthermore, while mice with impaired ability to create obesogenic immune memory have similar metabolic responses as wildtype mice to stable obesity, they are protected from the worsened glucose tolerance associated with weight cycling. Our data are the first to target metabolic consequences of weight cycling through an immunomodulatory mechanism. Thus, we propose a new avenue of therapeutic intervention by which targeting memory T cells can be leveraged to minimize the adverse consequences of weight cycling. These findings are particularly timely given the increasing use of efficacious weight loss drugs, which will likely lead to more instances of human weight cycling.

## Introduction

Obesity is increasingly prevalent worldwide and serves as a catalyst for a multitude of diseases including cardiovascular disease, diabetes, and cancer. Weight loss is encouraged for individuals with obesity (1); however, it is extremely challenging and only about 20% of individuals are successful in maintaining weight loss for an extended period of time (2). Instead, most individuals experience weight cycling, defined as bouts of weight loss and weight regain, for a significant portion of their lives. Compared with stable obesity, weight cycling further exacerbates metabolic disease (3–5). Despite the success of contemporary weight loss medicines, adherence to these drugs is <50% after a year of treatment (6–9), resulting in rapid weight regain. These clinical and real-world findings of weight rebound underscore the impetus for unraveling the mechanisms by which weight cycling increases metabolic disease risk.

Many of the diseases associated with obesity are a consequence of increased inflammation and immune cell infiltration into metabolic tissues, primarily adipose tissue (10). The adipose tissue environment is characterized by a collection of both innate and adaptive immune cells, the numbers and profiles of which dynamically change in response to obesity (11). Lean adipose tissue contains anti-inflammatory immune cells such as tissue resident macrophages, regulatory T cells (Treg) and T helper (Th2) cells, all of which contribute to maintenance of adipose tissue homeostasis. However, obesity perturbs the immune cell balance and leads to the accumulation of pro-inflammatory immune cells such as lipid associated macrophages and CD8^+^ T cells. This hyperinflammatory state promotes the development of insulin resistance and systemic glucose dysregulation (12). CD8^+^ T cells are implicated in obesity-induced insulin resistance (13), and conserved T cell receptor (TCR) sequences have been identified on clonally expanded CD8 ^+^ T cells in obese adipose tissue (14–17). Thus, there is strong evidence supporting the concept of “obesogenic memory” (15, 16, 18–21). While much work has characterized the immune cell niche in obese adipose tissue, less is known about the immune cell profile of adipose tissue in the context of weight loss and weight cycling. Glucose tolerance and insulin sensitivity improve, yet the proinflammatory immune profile in the adipose tissue remains after weight loss (10, 18, 22, 23). Others have reported that previously obese mice regain weight at a faster rate, display greater fat accumulation, and acquire an adaptive immune response in adipose tissue (18, 24–26). The recovery towards a normalized immune phenotype in adipose tissue after weight loss may contribute to worsened glucose tolerance associated with weight regain (22, 27, 28).

T cell memory, a crucial facet of adaptive immunity, involves the ability of specialized T cells to "remember" previously encountered pathogens to facilitate a rapid and more effective immune response upon antigenic re-exposure. One costimulatory interaction that promotes T cell memory formation is the CD27 receptor/CD70 ligand axis (29). Low levels of CD27 are constitutively expressed on NK cells, memory B cells, and quiescent T cells. Binding of the co-stimulatory ligand CD70 (found on antigen presenting cells; APCs) to CD27 (on naïve CD4^+^ and CD8^+^ T cells) leads to T cell proliferation, activation, and memory formation (30). Obesity significantly increases the abundance of memory T cells within adipose tissue (16, 31, 32), suggesting that the CD27/CD70 axis may be a a new therapeutic target that can be leveraged in metabolic disease.

We tested the hypothesis that T cell memory is causative in the exaggerated metabolic dysfunction resulting from weight cycling. We report that mice lacking CD70 (*CD70^-/-^*) become glucose intolerant in a stable obesity setting but are protected from worsened glucose tolerance associated with weight cycling. Additionally, the clonal expansion of adipose tissue T-cells is significantly reduced in *CD70^-/-^* mice. Thus far, most interventions that combat the effects of weight cycling take a surgical or pharmacological approach. This is the first time the metabolic consequences of weight cycling have been addressed through an immunomodulatory lens. It is becoming clearer that the complexity of obesity requires a multipronged care strategy. Here we suggest that targeting immune memory is an important component in improving metabolic function.

## Results

### CD8^+^ T cells are clonally enriched in adipose tissue during weight loss and weight cycling

To assess the impact of weight loss and weight cycling on T cell memory formation in adipose tissue, we performed analysis on single cell RNAseq (scRNAseq) data (22) from mice that had undergone 9-week cycles of high fat diet (HFD) and low-fat diet (LFD) feeding to create models of obesity, weight loss (WL), and weight cycling (WC) as previously reported (22, 27). At the end of the study, the obese and WC groups had the same body weight and adiposity as did the lean and WL groups (22). Droplet-based scRNAseq coupled with CITEseq was conducted on CD45^+^ cells extracted from the epididymal adipose tissue of four mice from each of the diet groups. A total of 33,322 single cells passed quality control and were categorized into 32 unique clusters (Fig. 1A-B). Six unique T cell subclusters were identified in the single cell dataset (Fig. 1C-D). A gene signature of T cell memory (33) is also enriched in the CD8^+^ T cells from obese, WL, and WC mice as compared with lean controls (Fig. 1E).

**Figure 1:**
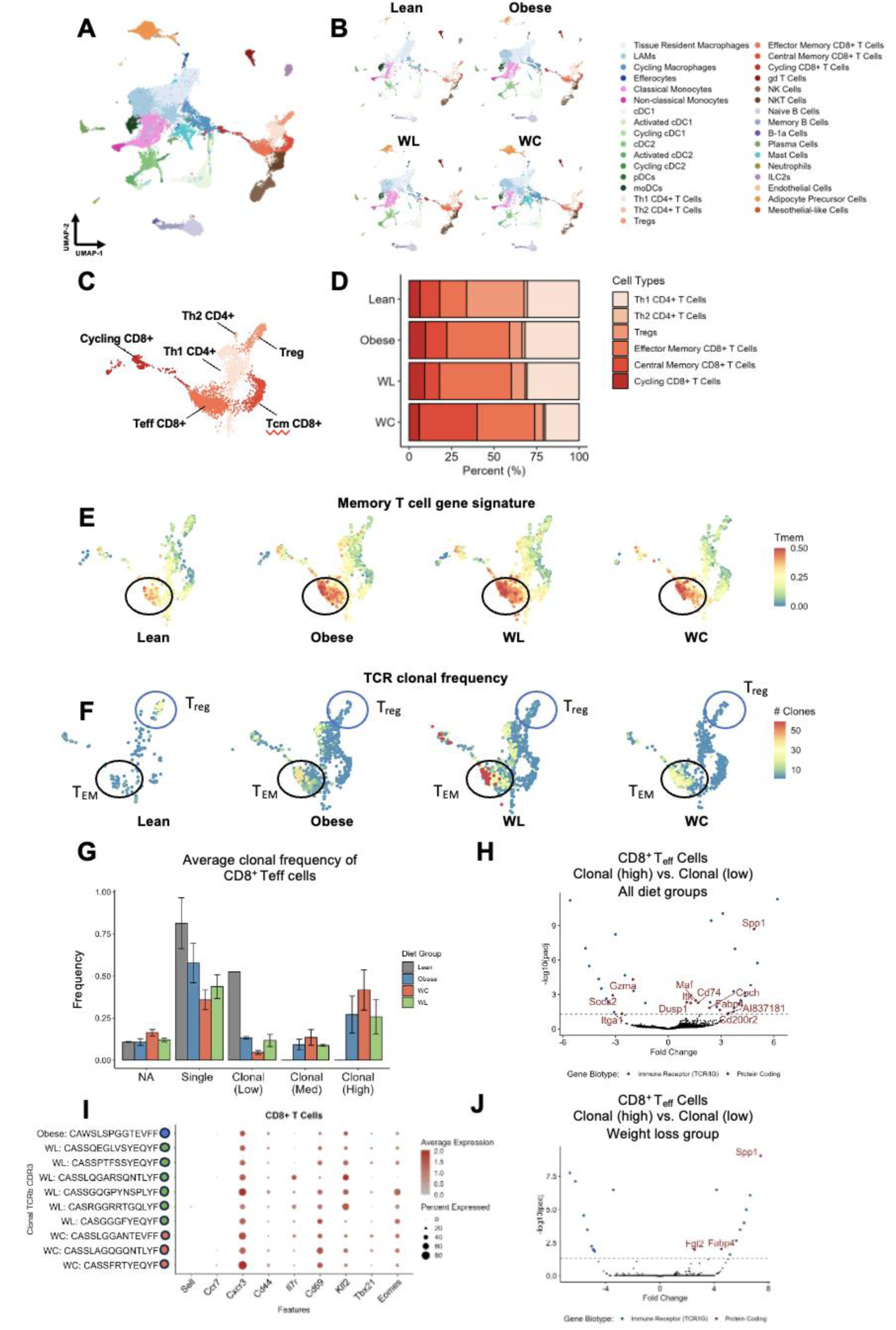
CD8^+^ memory T cells are clonally expanded in obese adipose tissue. A. Unbiased clustering of 24,156 single cells labeled broadly by cell type category and colored by high-resolution cell type identities via Uniform Manifold Approximation and Projection (UMAP). Populations include lipid-associated macrophages (LAMs), conventional dendritic cells (cDCs), monocyte-derived DCs (moDCs), T helper (Th) cells, T regulatory cells (T_regs_), gamma-delta (gd) T cells, and natural killer (NK) cells. B. UMAP by diet group for lean, obese, weight loss, and weight cycled mice. C. T cell populations identified. D. Proportion of T cell populations by diet group for lean, obese, weight loss, and weight cycled mice. E. Memory gene expression profile in T cell populations by diet group for lean, obese, weight loss, and weight cycled mice. F. Clonally expanded T cell populations by diet group for lean, obese, weight loss, and weight cycled mice. Black circles denote CD8^+^ T cells, blue circles denote T_regs_. G. Clonal frequency in CD8^+^ T cell populations by diet group for lean, obese, weight loss, and weight cycled mice. H. Differential expression of high clonal vs. low clonal CD8^+^ T cells across all diet groups. I. Averaged expression of memory T cell genes in the top 10 most enriched adipose tissue CD8^+^ T cell clonotypes organized by diet group in which they were identified. J. Differential expression of high clonal vs. low clonal CD8^+^ T cells across the weight loss group only. WL, weight loss; WC, weight cycled.

V(D)J sequencing was performed on scRNAseq samples to recover paired α and β TCR chain information to determine the effects of weight loss and weight cycling on T cell clonality. To visualize clonal enrichment, cells were binned into groups based on their clonal size (i.e. the number of cells with a matching clone). Clonal bins were determined as either single clones (only one cell with a given clonotype, i.e. not clonal) or into one of two bins (low clonal, 2-6 cells with the same TCR clonotype or high clonal, >6 cells with the same clonotype). Immunosuppressive T_regs_ had reduced clonality in obese, WL, and WC mice compared to lean mice (Fig. 1F). This lean-to-obese loss of T_reg_ TCR clonality is consistent with published work (34), but the observation that T_reg_ clonality is not recovered after weight loss is novel. There was an increase in CD8^+^ T_eff_ memory clonality in obese compared to lean animals that has been published by our group (17) and others (15, 34, 35). Data here shows that CD8^+^ T_eff_ memory clonality is not only retained in WL and WC mice, but is further increased in these groups (Fig. 1G). Analysis of the differential expression profile of high compared to low clonal cells across all diet groups indicates an increase in genes associated with immune cell activation and cytokine production (Fig. 1H). Additionally, an evaluation of the highly expressed clonotype sequences amongst the effector memory CD8^+^ T cell population show that 60% of the top clones are from WL animals. These highly clonal cells also show clear expression of T cell memory genes such as *Cxcr3*, *CD44*, and *Cd69* (Fig. 1I). Differential expression of the high versus low clonal cells in the WL group shows upregulated expression of *Spp1* (osteopontin), *Fgl2* (fibrinogen like protein 2), and *Fabp4* (fatty acid binding protein 4) (Fig. 1J). Taken together, these data suggest a role for obesogenic memory in the adipose tissue of WL and WC mice potentially contributing to metabolic dysfunction.

### *CD70*^-/-^ animals are protected from worsened glucose tolerance due to weight cycling

CD8^+^ T cell memory formation is guided by CD27/CD70 co-stimulatory interaction. Upon antigenic stimulation, CD70 is upregulated on APCs resulting in activation of CD27 signaling on CD8 ^+^ T cells, which promotes immune memory and broadens TCR repertoire (36). The CD27/CD70 axis increases the number of effector T cells and contributes to memory formation by counteracting T cell death during the contraction phase of adaptive immunity. Thus, we next leveraged the memory formation function of the CD27/CD70 axis to test whether interruption of this interaction prevents worsened metabolic outcomes with weight cycling. We first tested whether CD70 is implicated in metabolic dysfunction assocated with diet induced stable obesity. Eight-week-old male and female *CD70^-/-^* and littermate control wild-type (WT) mice were placed on either 10% low fat diet (LFD) or 60% high fat diet (HFD) for 16 weeks (Supplementary Fig. 1A). In both male and female groups, HFD increased body mass (Supplementary Fig. 1B-C). Glucose tolerance was assessed at weeks 8 and 16 via an intraperitoneal glucose tolerance test (GTT). Obese mice showed impaired glucose tolerance compared to lean mice, however there were no differences by genotype (Supplementary Figure 1D-E). These data suggest that in the absence of a secondary stimulus (weight regain), the CD70 associated T cell memory phenotype is inconsequential for glucose tolerance.

Next, littermate *CD70^-/-^* and WT male mice were placed on a 27-week weight cycling diet paradigm (Fig. 2A) to test whether interruption of the CD27/CD70 axis during weight regain ameliorates metabolic dysfunction. Change in body mass, weekly energy intake, and cumulative energy intake were not significantly different by genotype between obese and WC groups or between the lean and WL groups (Fig. 2B, Supplementary Fig. 2A-C). Fat mass was lower in *CD70^-/-^* versus WT mice in obese and WC conditions (Fig. 2C-D). No differences in fat mass were shown between WT and *CD70^-/-^* mice in lean or weight loss groups. Lean *CD70^-/-^* mice had greater lean mass than WT controls, whereas no differneces in lean mass were found between genotype within obese, WC, or weight loss groups, respectively. Mice that consumed high fat diet had increased fasting glucose as expected (Fig 2E). An oral GTT was performed before each diet switch and at the end of the study. At weeks 9 and 18 there were no significant differences in glucose tolerance by genotype fro any of the diet groups (Supplementary Fig. 3A-B). Similarly, at week 27, there were no significant differences in glucose tolerance in the lean, obese, and WL groups when comparing WT and *CD70^-/-^* (Fig. 2F-G, I). Remarkably, only in the WC groups, loss of CD70 mitigated the worsened glucose tolerance observed in WT mice (Fig. 2E & I). These data support that T cell memory is causative in exacerbating glucose intolerance with weight cycling in males.

**Figure 2:**
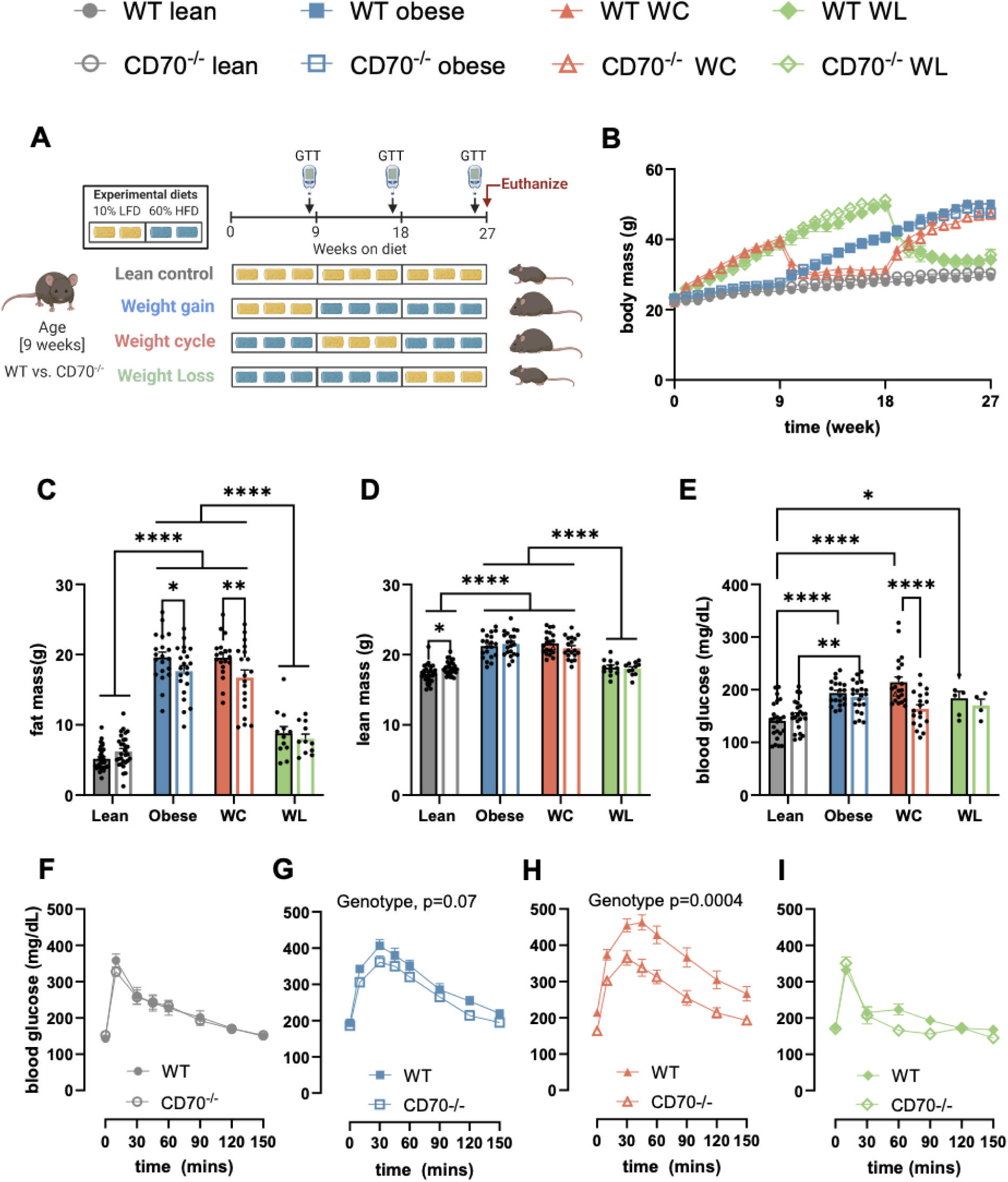
Weight cycling induced worsened glucose tolerance is corrected by the loss of CD70 in male C57BL/6J mice. A. Male littermate C57BL/6J mice were fed LFD or HFD for indicated 9- week blocks over a 27-week period. B. Weekly body weight over the course of 27 weeks. C-D. Final fat mass and lean mass at the 27-week end point via echo MRI. E. Fasting blood glucose from tail collected at 26-week time point. F-I. Oral glucose tolerance test at week 26 of diet feeding. A glucose bolus (3.5 g/kg fat free mass gavage) was delivered after 5 hours of fasting during the light cycle. Two-way repeated measures ANOVA with time and genotype (WT vs. CD70^-/-^) as factors were conducted. Multiple comparisons were assessed with Tukey *post hoc* testing. Data are means ± SE of 4-30 mice/group. WL, weight loss; WC, weight cycled. Graphics were generated via BioRender (BioRender.com). ** p=0.009 ***p=0.0005 ****p<0.0001

Female WT and *CD70^-/-^* animals were also placed on the same 27-week diet paradigm as their male littermates. Change in body mass and weekly energy intake were not significantly different by genotype between obese and weight cycled groups (Supplementary Figure 4A-B). Weight gain in female mice is driven by an increase in fat mass specifically (Supplementary Figure 4C-D). In contrast to their male littermates, there were no significant differences in fasting glucose or glucose clearance between WT and *CD70^-/-^* for any diet group (Supplementary Figure 4E-H).

### Loss of CD70 blunts the increase of adipose tissue CD8^+^ T cells

Previous work from our lab has shown that weight cycling in WT mice drastically changes the immune cell milieu in adipose tissue. To determine if differences in immune populations contribute to the improved glucose tolerance observed in *CD70*^-/-^ WC mice, we next profiled the immune environment using droplet-based scRNAseq on isolated CD45^+^ cells as previously described (22). We compared immune profiles of epididymal adipose tissue depots from animals according to the paradigm depicted in Figure 2A. We selected 4 representative animals per diet/genotype group. Across the four sample groups (32 mice), a total of 154,816 single cells were identified and fell within 32 unique clusters (Fig. 3A). We did not observe any significant difference in the proportion of macrophage or dendritic cell subpopulations between WT and *CD70^-/-^* across all four diet conditions (Fig. 3B-C). However, when comparing the T cell subpopulations, we observed that *CD70^-/-^* adipose tissue had significantly more CD4^+^ Th1 cells at the expense of effector memory CD8^+^ T cells in the obese, WL, and WC groups (Fig. 3D). These results are especially interesting in the context of weight loss because these mice are the same weight as their lean littermates. Differential abundance analysis of WL mice further highlights that neighborhoods (defined by cells with similar transcriptional patterns) of T and B cells are decreased in *CD70^-/-^* adipose tissue while stromal and plasma cells are enriched (Fig. 3E). These data further implicate a role for obesogenic memory in the context of diet-induced obesity and WL.

**Figure 3:**
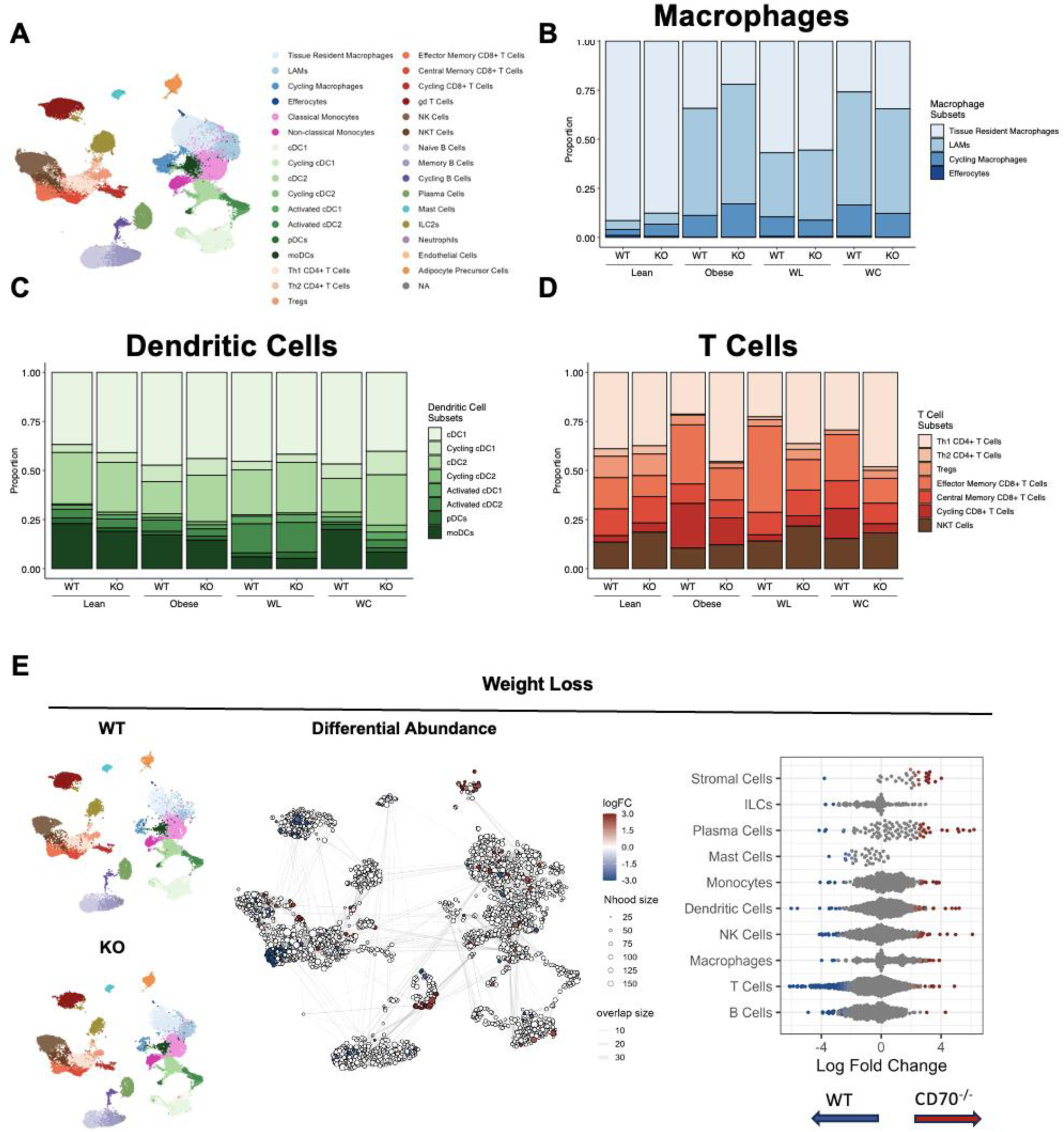
Loss of CD70 alters adipose tissue T cell populations across diet groups. A. Unbiased clustering of 154,816 single cells labeled broadly by cell type category and colored by high- resolution cell type identities via Uniform Manifold Approximation and Projection (UMAP). Populations include lipid-associated macrophages (LAMs), conventional dendritic cells (cDCs), monocyte-derived DCs (moDCs), T helper (Th) cells, T regulatory cells (T_regs_), gamma-delta (gd) T cells, and natural killer (NK) cells from WT and CD70^-/-^ adipose tissue. B. Macrophage proportions across diet and genotype groups. C. Dendritic cell proportions across diet and genotype groups. D. T cell proportions across diet and genotype groups. E. Differential abundance amongst 10 cell types between WT and CD70^-/-^ WL adipose tissue. WL, weight loss; WC, weight cycled.

### Loss of CD70 prevents clonal expansion of CD8^+^ T cells in weight loss

Because memory formation is likely to persist during the weight loss phase of our diet paradigm, coupled with the strong T cell memory phenotype observed in WL adipose tissue (Figure 3D), we next focused in on the weight loss condition. A second cohort of WL WT and *CD70^-/-^* mice were generated. To assess the role of T cell memory formation in *CD70^-/-^* WL mice, the T cell populations in the epididymal adipose tissue from WL animals were analyzed. To obtain enough T cells, CD4^+^/CD8^+^ were first sorted and 20,805 T cells containing immune receptors were assessed for clonal analysis (Fig 4A). Single cell sequencing of sorted T cells from WL mice of both genotypes revealed a shift in the T cell populations found in the epididymal adipose tissue marked by an increase in Th1 CD4^+^ T cells and a decrease in the CD8^+^ T cells in the *CD70^-/-^*animals (Fig. 4B). We observed an overall decrease in T cell clonality in the *CD70^-/-^* as compared to WT mice (Fig. 4C). To further interrogate these changes in clonality, cells were binned into one of three categories according to clonal size (single or nonclonal, low clonal < 56 clones, high clonal > 56 clones). In WT mice, the low clonal and high clonal populations were primarily effector memory CD8^+^ T cells (Fig. 4D-F), consistent with our previous findings (Figure 1). In contrast *CD70^-/-^* mice display a shift from clonal dominance in the CD8^+^ T cell population to clonal expansion in Th1 CD4^+^ cells (Fig. 4D-F).

**Figure 4:**
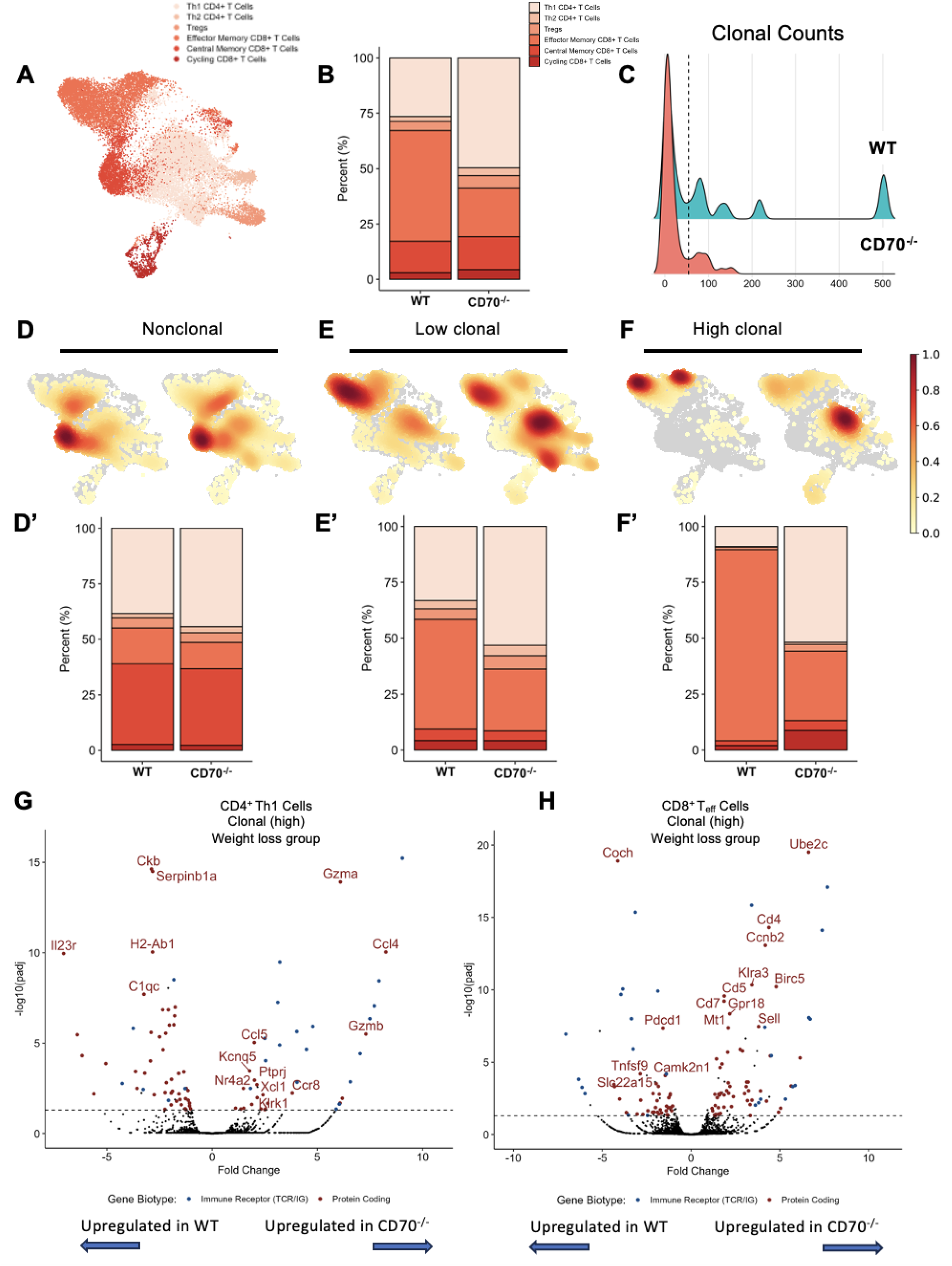
Loss of CD70 blunts the increase in adipose tissue CD8+ T cells and clonal expansion during weight loss. A. Proportion of T cell populations by genotype in weight loss adipose tissue. B. Clonal T cell abundance by genotype in weight loss adipose tissue. C-E. Density plot showing which T cell subsets are most clonally expanded by genotype in weight loss adipose tissue. C’- E’. Proportion of T cell subsets in highly clonal groups by genotype in weight loss adipose tissue. F. Differential expression of highly clonal CD4+ Th1 cells in CD70-/- vs. WT from weight loss adipose tissue. G. Differential expression of highly clonal CD8+ Teff cells in CD70-/- vs. WT from weight loss adipose tissue.

Differential expression amongst the highly clonal Th1 CD4^+^ cells isolated from adipose tissue of WL mice shows an increase in cytotoxic and chemoattractant genes (Fig. 4G). Differential expression amongst CD8^+^ T cells under the same conditions shows an increase in cell cycle and proliferation genes (Fig. 4H). Taken together, the shift in immune populations observed in WL adipose tissue results suggest that CD70 is a critical player in adipose tissue immune memory formation. The shift towards a highly clonal Th1 CD4^+^ population and away from CD8^+^ effector T cells may contribute to the metabolic protection observed in *CD70^-/-^* WC animals.

## Discussion

The adaptive immune system protects against complex and evolving pathogens and also recognizes danger signals from damaged or mutated cells (37). Modern environments pose different adaptive challenges to the immune system, contributed to by positive energy balance due to increased consumption of calories and low levels of physical activity. Thus, our current lifestyle, characterized by accumulated ‘metabolic stress’ can lead to an overactive or dysregulated immune response, results in negative health consequences. Herein, we make the case that a secondary challenge with HFD after weight loss incites a metabolic stress that is met with augmented and sustained memory T cell responses in adipose tissue. The result is worsened glucose tolerance and tonic elevation in inflammation. Using a mouse model of diet-induced weight cycling we show that: 1) CD8^+^ T cells in adipose tissue of WT mice display increased clonality in obesity, weight loss, and weight cycling; 2) inhibition of T cell memory formation constrains clonal expansion of CD8^+^ T cells in adipose tissue; and 3) loss of obesogenic memory abrogates the metabolic consequences of weight cycling but not stable obesity.

Our previous work has shown that weight cycling causes worsened glucose tolerance compared with obesity alone (22, 27, 38). Additionally, many groups have shown that clonality of CD8^+^ T cells is increased in adipose tissue, suggesting that CD8^+^ T cells in this tissue are primed by specific antigens. This increase in clonality is not only retained in WL and WC mice, it is further increased. This is particularly interesting for the WL group as these mice are metabolically indistinguishable from lean counterparts. *CD70^-/-^* WC animals demonstrate reduced CD8^+^ T cell clonality as compared to WT animals with a shift towards Th1 CD4^+^ T cells in their adipose tissue. While Th1 cells are generally associated with inflammation, they can also play a regulatory role in balancing the activity of other T cell subsets, such as Th2 cells and T_regs_ (31). The balance between pro-inflammatory Th1 cells and anti-inflammatory T_regs_ in adipose tissue can determine the overall immune environment and influence the level of inflammation in the tissue. The observation that *CD70^-/-^* mice are protected only from the worsened metabolic consequences of weight cycling suggests a role for CD70 in the development of obesogenic memory, and more importantly, that inhibiting CD70 may confer some benefits for individuals who weight cycle. This is one of the first examples in which blocking a component of T cell memory formation has proven to be beneficial in the context of metabolic disease. This has many important implications both at the level of the adipose tissue and potentially for other organ systems controlling glucose tolerance (e.g., liver, pancreas, skeletal muscle). CD70 has already emerged as a promising therapeutic target in human cancers due to its restricted expression in normal tissues but significant overexpression in a variety of malignancies, including renal cell carcinoma, glioblastoma, and certain hematological cancers like lymphoma. Targeting CD70 with monoclonal antibodies, antibody-drug conjugates, or other immunotherapeutic strategies has shown potential to selectively eliminate CD70-expressing cancer cells while sparing healthy tissues, making it a highly attractive candidate for cancer treatment (39, 40). Taken together, our data reveals a potential new target for CD70 therapies in human adipose tissue.

Losing excess weight undoubtedly improves metabolic health (41), but sustained weight loss remains challenging due to strong neuroendocrine drive to regain weight. This is particularly concerning because weight loss alone does not erase the inflammatory signature in adipose tissue (42, 43) and this inflammation worsens with weight regain (22, 44). In response to rising obesity rates, weight loss drugs have gained global adaptation. Since their approval for diabetes treatment in the mid-2000s, incretin-based drugs have become some of the most prescribed in the United States for weight loss (45). However, when patients discontinue use, they regain a significant portion of the weight lost, often leading to further weight regain (i.e. weight cycling). This is troubling given recent discoveries, including findings from this study, that weight regain worsens health outcomes (5, 46, 47). Intriguingly, incretin-based therapies may impact the inflammatory environment of the adipose tissue thereby influencing the activity and function of immune cells (48) and this may be independent of weight loss. These results raise the possibility that these therapies might have broader effects on immune regulation, but specific research on how they impact memory T cells within adipose tissue is not known. None-the-less, targeting the adaptive immune system may help mitigate some of the adverse consequences of weight regain.

While modern interventions such as bariatric surgery and weight loss drugs have garnered significant attention, it is important to recognize that caloric restriction has long been associated with metabolic health improvements and may offer additional benefits in mitigating the adverse effects of weight cycling. Studies have shown that caloric restriction, defined as negative energy balance without malnutrition, can protect tissues from chronic inflammation by reducing pro-inflammatory cytokines, enhancing immune regulation, and improving mitochondrial function (49, 50). Importantly, caloric restriction also helps safeguard the adaptive immune system by preventing the overactivation of immune cells, such as T cells, which can become dysregulated in the presence of chronic metabolic stress. By promoting the activity of T_regs_ and limiting the expansion of memory T cells associated with inflammation, caloric restriction helps maintain a controlled immune response (51). These effects reduce the immune system’s burden, preserving its ability to respond appropriately to pathogens without contributing to the chronic low-grade inflammation seen in obesity. Thus, caloric restriction may complement other therapies and help prevent the metabolic stress and immune dysfunction that follow weight regain.

Robust obesogenic memory formation is not observed in *CD70^-/-^*mice suggesting weak costimulation or insufficient antigen presentation. Differential expression analysis of highly clonal WL cells showed that *CD70^-/-^*cells have a strong gene signature for cell proliferation, migration, and chemoattraction relative to WT. This suggests that in the absence of CD70, T cells activate alternative gene programs, though they lack the full range of tools needed to effectively combat the chronic inflammatory state. This shift towards effector Th1 CD4^+^ T cells highlights the necessity of CD70 for proper memory formation.

Antigen presentation is a critical step in both obesity-associated inflammation and memory formation, bridging the innate and adaptive immune systems that drive the metabolic defects associated with weight cycling. Multiple cell types in adipose tissue, including macrophages, dendritic cells, B cells, and adipocytes, can function as APCs (52–55). Certain APCs have been shown to develop obesogenic memory themselves, though through mechanisms distinct from those observed in the adaptive immune system. We have previously shown a form of innate immune memory in adipose tissue macrophages (28). Memory formation in adipocytes is due to epigenetic modifications in adipocytes (21, 28), offering another therapeutic avenue to combat obesity. Because our studies utilized global *CD70^-/-^* mice, the specific APC type responsible for the reduction of immune memory and amelioration of worsened metabolic consequences of weight cycling cannot be identified. In addition, the antigen or class of antigens being presented to T cells has yet to be identified. Our previous work studying the complementarity determining region 3 of the T cell receptor of clonal CD8^+^ T cells suggests that modification of proteins via negatively charged, highly reactive γ-ketoaldehydes called isolevuglandins, may be a family of candidates(17). More recent data highlight Cd1d expressing cells that present endogenous lipid antigens to natural killer T (NKT) cells, influencing inflammation and metabolic processes in obesity (56–59). The pro-lipid environment in obese adipose tissue facilitates the presentation of lipid-enriched antigens to T cells inducing T cell memory. An opportunity exists to further investigate this lipid- T cell interaction. Our data show that preserving an anti-inflammatory, clonally diverse immune population within adipose tissue may mitigate some of the negative health consequences of weight cycling.

Much remains to be explored regarding the temporal window in which T cell memory formation is crucial for the metabolic benefits we have observed. In using whole body knockouts, our current studies are unable to answer this question. Future studies are needed to investigate whether depleting CD70 during the early phase of the weight loss period (weeks 9-12 of diet) or the early phase of the weight regain (weeks 18-21 of diet) better confer improved glucose tolerance. Alternatively, APC-specific CD70 knockout mice could be used to better address this question as whole tissue metabolism may be impacted by cell depletion alone or the altered phenotype of the immune cell.

Our present findings demonstrate that blunting obesogenic memory effectively mitigates the metabolic consequences of weight cycling, underscoring the critical role of immune memory in obesity-related dysfunction. Given the complexity of obesity, a multipronged care strategy is essential for addressing its various physiological and metabolic effects. Here, we highlight targeting immune memory as a pivotal component for improving metabolic outcomes. Taken together, our findings support a novel pathway to reduce the detrimental effects of weight cycling, which may advance therapeutic strategies for obesity management.

## METHODS

### Ethics Statement

All procedures were approved in advance and carried out in compliance with the Vanderbilt University Institutional Animal Care and Use Committee. Vanderbilt University is accredited by the Association for Assessment and Accreditation of Laboratory Animal Care International.

### Animals and Experimental Design

CD70 mice were a kind gift from the Ross Kedl laboratory at University of Colorado Anschutz Medical Campus on the C57BL/6N background and were backcrossed 10 generations onto the C57BL/6J background in our laboratory. Male C57BL/6J heterozygous mice were bred to generate wild-type and *CD70^-/-^* mice. Littermate controls were used in the studies documented in this manuscript. Wild-type and *CD70^-/-^* mice were placed into one of four diet groups at 8 weeks of age: lean, obese, weight cycled, or weight loss. Mice were maintained on their diet for 27 weeks, divided into 3 9-week cycles. Lean mice were maintained on low fat diet (LFD: 10%, Research Diets #D12450B, 3.85 kcal/g) for a total of 27 wks. Obese mice were maintained on LFD for the first 9 wk cycle and then transitioned to high fat diet (HFD: 60% fat, Research Diets #D12492, 5.24 kcal/g food) for the remaining 18 wks. Weight cycled mice were maintained on HFD for the first 9 wks, transitioned to LFD for the second 9 wks, and switched back to HFD for the final 9 wks. Weight loss animals were on HFD for 18 wks and then switched to LFD for 9 wks.

### Glucose tolerance testing

Glucose tolerance tests were performed by intraperitoneal or oral delivery as indicated in figure legends. Body composition was assessed to determine glucose dosing to lean mass for the glucose tolerance tests. Mouse body fat and fat free mass were measured by a nuclear magnetic resonance whole body composition analyzer (Bruker Minispec). After a 5 h fast, basal blood glucose levels were measured (0 min) followed by intraperitoneal injection of 2 g/kg fat free mass body mass or oral gavage of 3.5 g dextrose/kg fat free mass. Blood glucose was sampled via tail cut via a 150 min time course after intraperitoneal injection or oral gavage using a hand-held glucometer (Bayer Contour Next EZ meter). Glucose area under curve from baseline was calculated using the trapezoidal rule.

### Adipose tissue SVF isolation

Mice were euthanized by isoflurane overdose and cervical dislocation followed by perfusion with 20 mL PBS through the left ventricle. Epididymal AT depots were excised, minced and digested in 6 mL of collagenase for 30 min at 37°C. For flow cytometry applications AT was digested in 2 mg/mL type II collagenase (Sigma C6885; St. Louis, MO) and for single cell applications AT was digested in 2 mg/mL type IV collagenase (Worthington LS004189; Lakewood, NJ, USA). Digested AT was then vortexed, filtered through 100 μm filters, lysed with ACK buffer and filtered through 35 μm filters as previously described (22).

### Single cell sequencing

Mice were euthanized and the SVF from the epididymal AT was isolated and sorted as described (22). The SVF was incubated with anti-mouse Fc Block (BD Biosciences) at 1:200 for 10 min. For animals in obese and WC groups, the CD45+ fraction of the epididymal AT was isolated for scRNAseq. Cells from each mouse were labeled with unique hashtag antibodies (1:200) (Biolegend TotalSeq-C) and anti-CD45 microbeads (10 μL/sample) (Miltenyi #130-110-608). Biological replicates were pooled and sorted on a manual Miltenyi OctoMACS Separator. CD45^+^ cells were then labeled for CITE-sequencing using TotalSeq-C antibodies (Biolegend) for cell surface markers to identify major cell types. All immunolabeling was completed at a 1:200 dilution for 20 min at 4°C in the dark. Next, cells were stained with 0.25 μg/mL DAPI for FACS sorting and DAPI^-^ viable cells were collected for downstream processing and sequencing. All samples from the obese and WC groups were submitted and processed for sequencing on the same day to minimize batch effects. Sample preparation was conducted using the 5’ assay for the 10X Chromium platform (10X Genomics) targeting 20,000 cells per diet group. In all, 50,000 reads per cell were targeted for PE-150 sequencing on an Illumina NovaSeq6000. Samples processing and sequencing were completed within the same run in the VANderbilt Technologies for Advanced Genomics core (VANTAGE).

Another round of single cell sequencing was performed on CD45^+^ cells isolated from the epididymal AT of lean and WL animals. The same procedure for isolating the SVF was followed as described above however, SVF was stained with CD45 (1:200) (Biolegend; clone 30-F11; catalogue #: 103137) antibody and 0.25 μg/mL DAPI. The live CD45+ cells were isolated by gating on the DAPI- CD45^+^ fraction on the 4-laser FACSAria III, Bioprotect IV Baker Hood Enclosure. Furthermore, in the WL adipose tissue, SVF was also stained with CD4^+^ (1:200) (Biolegend clone RM4-4; catalogue #: 116022) and CD8^+^ (1:200) (Biolegend clone 53-6.7; catalogue#: 100707) antibody. Following the DAPI^-^ CD45^+^ sort, sample was also sorted for DAPI^-^

, CD45^+^, CD8^+^, CD4^+^ cells. The Chromium Next GEM Single Cell 5’ HT v2 kit from 10X Genomics was implemented for both lean and WL samples and run on a NovaSeq 6000.

### Single cell RNAseq Data Processing

Reads from newly generated single cell RNAseq data were processed using CellRanger v7.0.1. For initial quality control of droplets, dropkick v1.2.7 (60) was performed. Then, dropkick score, mitochondrial RNA content, total number of UMI counts, and feature counts informed thresholding for each individual sample. All cells with > 20% mitochondrial content were filtered. Doublet detection was performed using scDblFinder (61) and cells identified as doublets were filtered prior to downstream analysis. For ambient RNA estimation and correction, the R package SoupX (62) v1.6.2 was used. Because SoupX requires clustering information to estimate the contamination fraction, each sample was independently processed using Seurat v5.1 (63) with the *NormalizeData, FindVariableFeatures, ScaleData, RunPCA, FindNeighbors, and FindClusters* functions^38^. SoupX adjusted count matrices were then loaded into Seurat. For downstream analyses requiring Anndata objects, scDIOR (64) was used for object conversion and analyses were performed in Python v3.11 using Scanpy (65)v1.9.1.

### Dimensional Reduction and Data Integration

Linear dimensional reduction was performed using principal component analysis with the *RunPCA* function based on 3,000 highly variable genes detected by the *FindVariableFeatures* function. Thirty principal components were used for data integration. Concatenated data was integrated with Harmony (66). Linear dimensional reduction was performed using Uniform Manifold Approximation and Projection (UMAP) with the *RunUMAP* function using default settings.

### Reference Mapping and Label Transfer

Cells were annotated by mapping to a reference murine adipose tissue immune atlas (22). First, integration features were selected using the *SelectIntegrationFeatures* function and then transfer anchors were identified using the *FindTransferAnchors* function with 30 principal components in Seurat v5.1. Then, cell annotation labels were transferred using the *TransferData* function.

### Differential Abundance Analysis

Differential abundance testing was performed using MiloR v2.20 (67). Briefly, the Seurat object containing single cell RNAseq results was converted to a SingleCellExperiment (68)object. In MiloR, cell neighborhoods were identified using the following parameters: k = 24, d = 30, and prop = 0.02.

### Differential Expression Analysis

For differential expression testing, a pseudobulk approach was used in which counts for cells belonging to each independent sample were aggregated. Then, edgeR v4.4.1(69) was used to perform differential testing. Briefly, normalization factors were calculated using the *calcNormFactors* function. Differential testing was performed using the *glmQLFFit* and *glmQLFTest* functions. Significantly differentially expressed genes were determined by an adjusted p value < 0.05 and an absolute log_2_ fold-change > 1.

### Immune Receptor Data Processing

Filtered and annotated contig files from CellRanger v7.0.1 were imported into R and filtered for contigs with productive T cell receptors (TCRs). In instances where more than one TCR was observed for a cell barcode, the TCR with the most captured reads was retained. Sequenced TCR nucleotide data were imported into the Seurat object metadata for downstream plotting.

## Statistical Analyses

Student’s t-tests were run for between group comparisons. In experiments that contained more than two groups, one-way analysis of variance (ANOVA) or two-way ANOVA models were conducted with pairwise comparisons using Tukey or Sidak correction. Brown-Forsythe correction was applied to groups with unequal variance. Data are presented as mean ± standard error (SE). An adjusted p value of <0.05 was used to determine significance. Statistical analyses were performed using GraphPad Prism.

## Data Availability

Both the raw and processed scRNAseq and VDJ sequencing data in this study have been submitted to the NCBI Gene Expression Omnibus (GEO) and will be made available prior to publication. The GEO Accession Number is **GSE287246**.

## Author’s Information

AHH is now Vice-Provost and Senior Associate Dean for Faculty Affairs and Career Development at University of Texas Southwestern in Dallas, TX

## Acknowledgments

We acknowledge the following Vanderbilt University (VU) and Vanderbilt University Medical Center (VUMC) core facilities: VU Metabolic Mouse Phenotyping Center [VMMPC (NIH DK135073 (MMPC-Live); www.vmmpc.org) and DK020593 (DRTC)], Vanderbilt Flow Cytometry Shared Resource [Vanderbilt Ingram Cancer Center (P30CA068485)], and the Vanderbilt Digestive Disease Research Center (DK058404). scRNAseq and VDJ sequencing was performed in the Vanderbilt Technologies for Advanced Genomics (VANTAGE) core laboratory, which is supported in part by Clinical and Translational Science Award Grant 5UL1 RR024975-03, Vanderbilt Ingram Cancer Center Grant P30 CA68485, Vanderbilt Vision Center Grant P30 EY08126, and National Institutes of Health/National Center for Research Resources Grant G20 RR030956.

