## Supplemental Figures for "Interrupting T cell memory ameliorates exaggerated metabolic response to weight cycling"

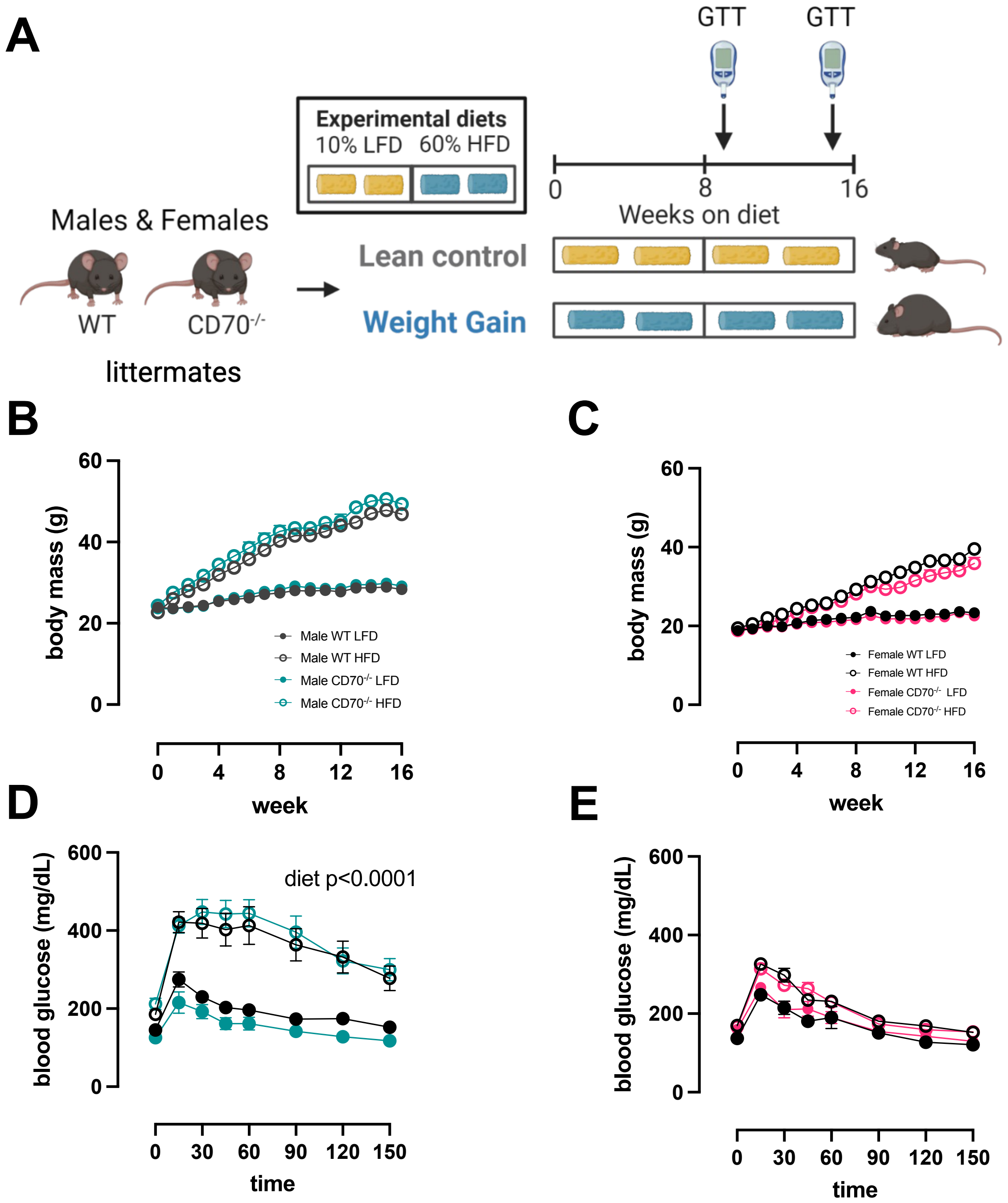


**Supplemental Figure 1:** Loss of CD70 does not impact weight gain in C57BL/6J males or females. A. Male and female littermate C57BL/6J mice were fed LFD or HFD for a 16-week period. B-C. Weekly body weight over the course of 16 weeks for males and females respectively. D-E. An intraperitoneal glucose tolerance test was conducted at week 15 of diet feeding for males and females respectively. A glucose bolus (2.0 g/kg fat free mass) was delivered after 5 hours of fasting during the light cycle. Two-way repeated measures ANOVA with time and group (WT LFD/HFD vs. CD70-/- LFD/HFD) as factors were conducted. Multiple comparisons were assessed with Tukey post hoc testing. Data are means ± SE of 8-10 mice/group.

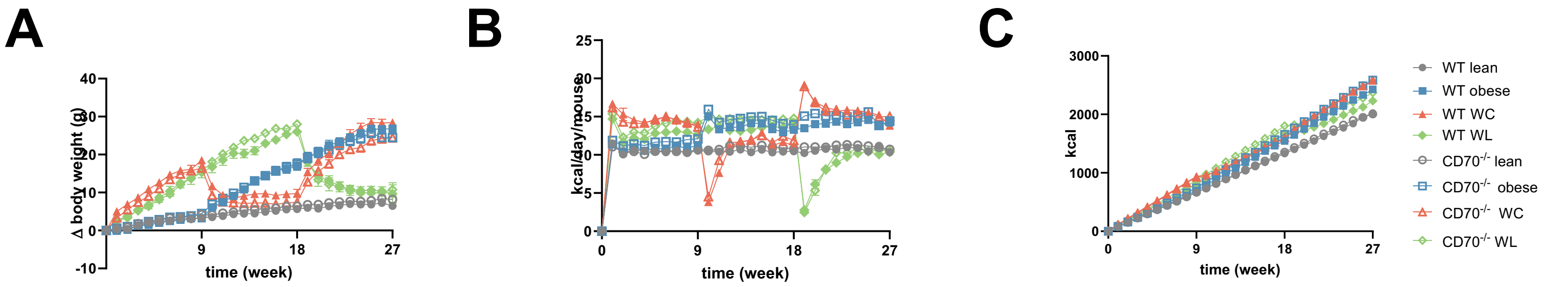


**Supplemental Figure 2:** Weight cycling induced worsened glucose tolerance is corrected by the loss of CD70 in male C57BL/6J mice. A. The change in body weight from week 0 to week 27. B. Daily caloric intake over the the 27-week study. C. Cumulative energy intake over the 27-week study. Data represent 10-30 mice/group.

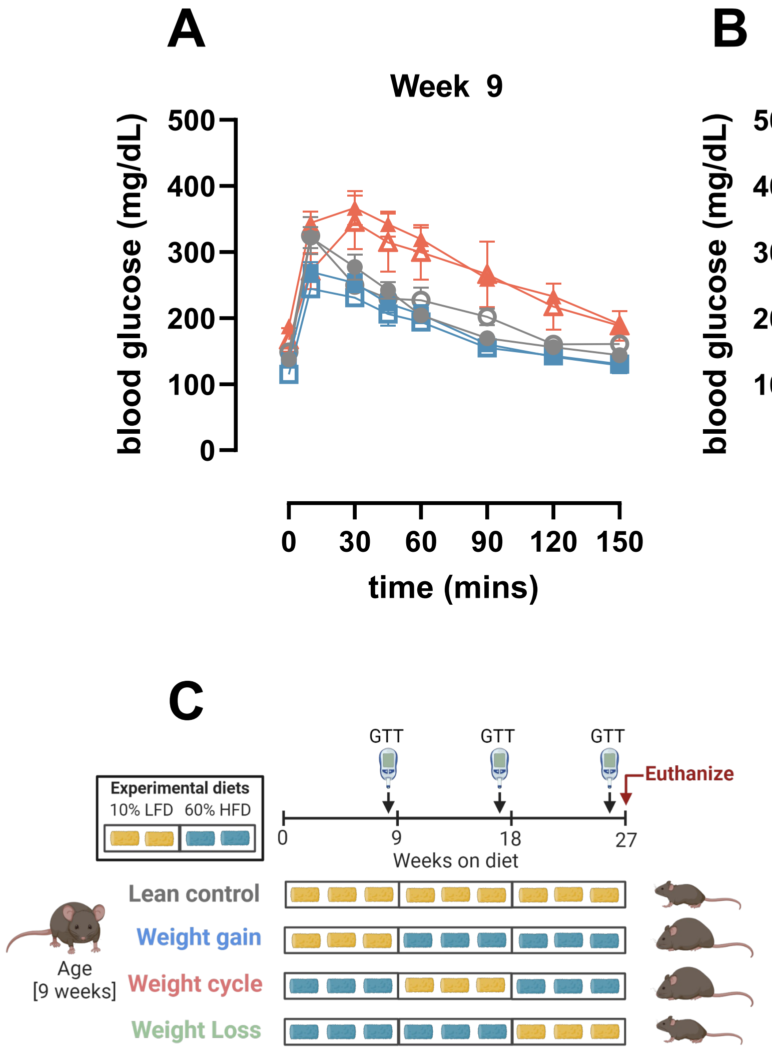


**Supplemental Figure 3:** Glucose tolerance of weight cycled males does not differ at week 9 and week 18. A. An oral glucose tolerance test was conducted at week 9 of diet feeding. B. An oral glucose tolerance test was conducted at week 18 of diet feeding. Two-way repeated measures ANOVA with diet and genotype as factors were conducted for panels A and B. Multiple comparisons were assessed with Tukey's *post hoc* testing. Data are means ± SE of 8-9 mice/group for panel A and 19-21 mice/group for panel B. C. Schematic showing when glucose tolerance tests were performed along the 27-week diet paradigm.

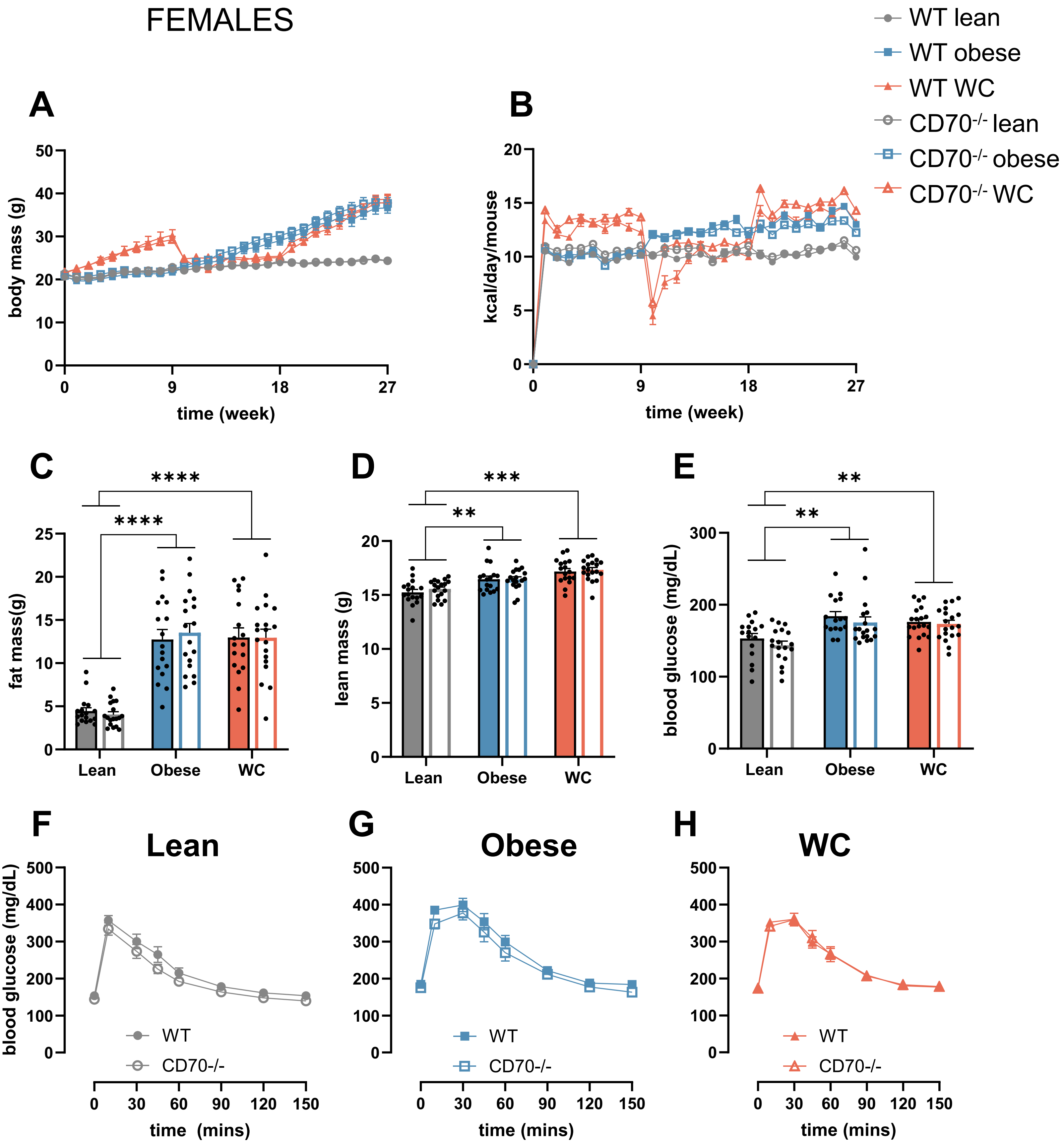


**Supplemental Figure 4:** Weight cycling does not exacerbate glucose tolerance and loss of CD70 does not improve glucose tolerance in C57BL/6J female mice. A. Weekly body weight over the course of 27 weeks. B. Daily caloric intake over the the 27-week study. C-D. Final fat mass and lean mass at the 27-week end point via echo MRI. E. Fasting blood glucose from tail collected at 26-week time period. F-H. A glucose tolerance test was conducted at week 26 of diet feeding. A glucose bolus (3.5g/kg fat free mass gavage) was delivered after 5 hours of fasting during the light cycle. The glucose excursion is reported. Two-way ANOVA with group and genotype (WT vs. CD70$^{-/-}$) as factors were conducted. Multiple comparisons were assessed with Tukey *post hoc* testing. Data are means ± SE of 17-20 mice/group. **p=0.01, ***p=0.001, ****p<0.0001.
